# Laser-facilitated epicutaneous immunization with SARS-CoV-2 spike protein induces ACE2 blocking antibodies in mice

**DOI:** 10.1101/2021.02.22.432259

**Authors:** Sandra Scheiblhofer, Stephan Drothler, Werner Braun, Reinhard Braun, Maximilian Boesch, Richard Weiss

## Abstract

The skin represents an attractive target tissue for vaccination against respiratory viruses such as SARS-CoV-2. Laser-facilitated epicutaneous immunization (EPI) has been established as a novel technology to overcome the skin barrier, which combines efficient delivery via micropores with an inherent adjuvant effect due to the release of danger-associated molecular patterns. Here we delivered the S1 subunit of the Spike protein of SARS-CoV-2 to the skin of BALB/c mice via laser-generated micropores with or without CpG-ODN1826 or the B subunit of heat-labile enterotoxin of *E.coli* (LT-B). EPI induced serum IgG titers of 1:3200 that could be boosted 5 to 10-fold by co-administration of LT-B and CpG, respectively. Sera were able to inhibit binding of the spike protein to its receptor ACE2. Our data indicate that delivery of recombinant spike protein via the skin may represent an alternative route for vaccines against Covid-19.

## Introduction

The spike protein has been identified as the target antigen of choice for SARS-CoV-2 vaccine development [1], pursued by numerous companies worldwide. Coronaviruses have a large, positive-sense single-stranded RNA genome and are enveloped, mostly expressing their characteristic spike protein as the only large envelope surface protein [2]. In case of the betacoronavirus SARS-CoV-2, the spike protein binds to its receptor angiotensin-converting enzyme 2 (ACE2) on host cells via the receptor binding domain (RBD) of its S1 subunit [3]. The spike S2 subunit facilitates endocytosis, eventually leading to fusion of viral and endosomal membranes and release of viral RNA into the cytoplasm [3, 4]. Antibodies that bind to the RBD have been shown to prevent its attachment to ACE2-expressing host cells, thereby neutralizing the virus [1, 5]. Immune responses during natural infections in humans are characterized by an initial wave of systemic IgG antibodies followed by a decline and formation of long-lived plasma cells. However, also the production of secretory IgA antibodies at mucosal surfaces has been reported [6, 7]. Compared to SARS-CoV, some structural differences within the RBD increase the ACE2 binding affinity of SARS-CoV-2, potentially explaining the higher transmission rate of SARS-CoV-2 [8].

Recently, immunization via the skin came into focus as an alternative to classical routes such as intramuscular or subcutaneous injections. Due to the high density of professional antigen presenting cells in the dermal and epidermal skin layers, vaccine uptake and antigen presentation can be improved, enabling dose reduction and providing increased efficacy in the elderly [9, 10]. We and others established generation of skin micropores via infrared laser beams as a painless and highly reproducible method to overcome the cornified outermost skin barrier and deliver large molecules such as biologicals [11–14]. This method has been recently tested in healthy human adults using epicutaneous patches containing recombinant pertussis toxin and was shown to induce recall antibody responses equivalent to those induced by injection of a commercial diphtheria-tetanus-acellular pertussis vaccine adjuvanted with alum [15].

In this proof-of-principle study, we investigated whether laser-facilitated epicutaneous immunization with recombinant SARS-CoV-2 S1 spike protein could induce antibody responses capable of preventing spike from ACE2 binding. For adjuvantation, we employed LT-B and CpG, as these two recently proved to be most effective in potentiating immune responses following skin-based immunization [16], and co-administered the universal T helper activating peptide PADRE [17].

## Materials and methods

### Reagents

The following reagents were used in the study: recombinant SARS-CoV-2 spike S1 protein (GenScript Cat. No. Z03501), TLR9 agonists CpG-ODN 1826 (Invivogen, Cat. No. TLR-1826), and LT-B (Sigma, Cat. No. E8656), PADRE peptide (AKFVAAWTLKAAA, Bachem AG, Bubendorf, Switzerland), EPIMMUN® skin patches (Pantec Biosolutions AG, Ruggell, Principality of Liechtenstein).

Adjuvants were diluted in endotoxin-free PBS to give the following stock solutions: LT-B was diluted to 10mg/mL, CpG-ODN 1826 was diluted to 5mg/mL, and PADRE was diluted to 5mg/mL. At this concentration, the latter was not fully soluble, resulting in a turbid suspension. Due to volume restrictions, this suspension was nevertheless used for immunization.

### Mice and their housing

Blood samples were drawn on days 0, 28, and 42 (experiment 1) and 0, 42, 63, and 84 (experiment 2) by puncture of the v. saphena.

Wild-type female BALB/c mice aged between six and eight weeks were purchased from Janvier Labs, France, and maintained at the animal facility of the University of Salzburg under SPF (specific pathogen-free) conditions according to FELASA guidelines. All animal experiments were conducted in compliance with EU Directive 2010/63/EU and have been approved by the Austrian Ministry of Education, Science and Research, under permit number GZ 2020-0.282.314. Mice were immunized at 8-10 weeks of age.

### Laser-assisted epicutaneous immunization

One day before laser microporation, the dorsal or ventral skin of the mice was depilated as previously described [16]. Epicutaneous immunization was performed with a P.L.E.A.S.E. ® Professional laser system (Pantec Biosolutions AG) using the following parameters: 0.7W, three pulses, total fluence of 8.3J/cm^2^, 9% pore density, 1cm diameter as previously described [16].

In the first experiment, mice were immunized with 25μg SARS-CoV-2 spike S1 protein with or without 50μg of PADRE peptide, 40μg LT-B, or a combination thereof. In the second experiment, mice were immunized with 25μg SARS-CoV-2 spike S1 protein with or without 40μg of CpG-ODN 1826. Formulations were applied to the laser-treated site in endotoxin-free PBS in a total volume of 29μL (the minimum volume we could achieve based on the concentrations of the stock solutions). Naïve mice (experiment 1) or laser-microporated mice sham treated with PBS (experiment 2) served as controls. Mice were kept on a heating pad until the solution had been completely taken up via the micropores. The treatment area was subsequently covered with an occlusive patch (EPIMMUN® skin patches), which was removed after 24h.

Mice were immunized three times at 14-day intervals (experiment 1) or 21-day intervals (experiment 2), and blood samples were taken 2 days before the first immunization and 12 days after the second and third immunization (experiment 1). In experiment 2, blood samples were taken 19 days after the second immunization, and 19 and 40 days after the third immunization.

At the time of sacrifice, bronchoalveolar lavage fluid (BALF) and splenocytes were harvested as previously described [14].

### Immunological assays

Serum and BALF samples were analysed by ELISA for SARS-CoV-2 specific IgG by a chemiluminescence-based ELISA as described [16].

The capacity of sera from immunized mice to inhibit the binding of spike protein to ACE2 was analysed using a SARS-CoV-2 Spike:ACE2 Inhibitor Screening Assay kit (BPS Biosciences, Cat. No. 79931) according to the manufacturer’s instructions. Briefly, soluble ACE2 receptor was preincubated with serial dilutions (1:5 to 1:15625) of sera from immunized mice and then added to ELISA plates coated with spike RBD protein. The amount of bound receptor was detected by a chemiluminescence reaction. As reference, serial dilutions of the spike S1 neutralizing antibody clone 414-1 (BPS Bioscience, Cat. No. 200916) were used. This antibody clone was isolated from a convalescent COVID-19 patient and showed the highest SARS-COV-2 neutralizing capacity [18].

Spike protein-specific T cell responses were detected by a standard ELISPOT assay using the matched clones 11B11 and BVD6-24G2 for IL-4, and XMG1.2 and R4-6A2 for IFN-γ. Briefly, 2 x 10^5^ splenocytes were incubated overnight in T cell medium (RPMI-1640, 10% FCS, 25 mM Hepes, 2 mM L-Glu, 100 μg/mL streptomycin, 100 U/mL penicillin) with or without 20μg/mL spike S1 protein. The assay was finally developed using AEC substrate (Sigma).

## Results

After the third immunization, mice immunized with the SARS-CoV-2 spike S1 protein displayed high titers of specific serum IgG in two independent experiments (Fig. 1A and B). LT-B significantly boosted IgG levels, whereas PADRE had a negative impact on immunogenicity (Fig. 1A). The latter observation was surprising and may have been associated with the precipitates present in the formulation. Mice immunized with CpG-ODN 1826 adjuvant showed the highest antibody titers of all groups (Fig. 1B). Increasing the interval between immunizations had no significant effect on the magnitude of the antibody response, as immunization with spike S1 protein in the absence of adjuvant (S1) resulted in basically the same titers in experiments 1 and 2 (Fig. 1C and D). CpG induced a faster onset of the immune response (Fig. 1D), even though this was solely tested with the longer immunization intervals. As shown in Fig. 1D, IgG titers remained stable after the third immunization and decreased only slightly between days 63 and 84.

**Fig. 1.**
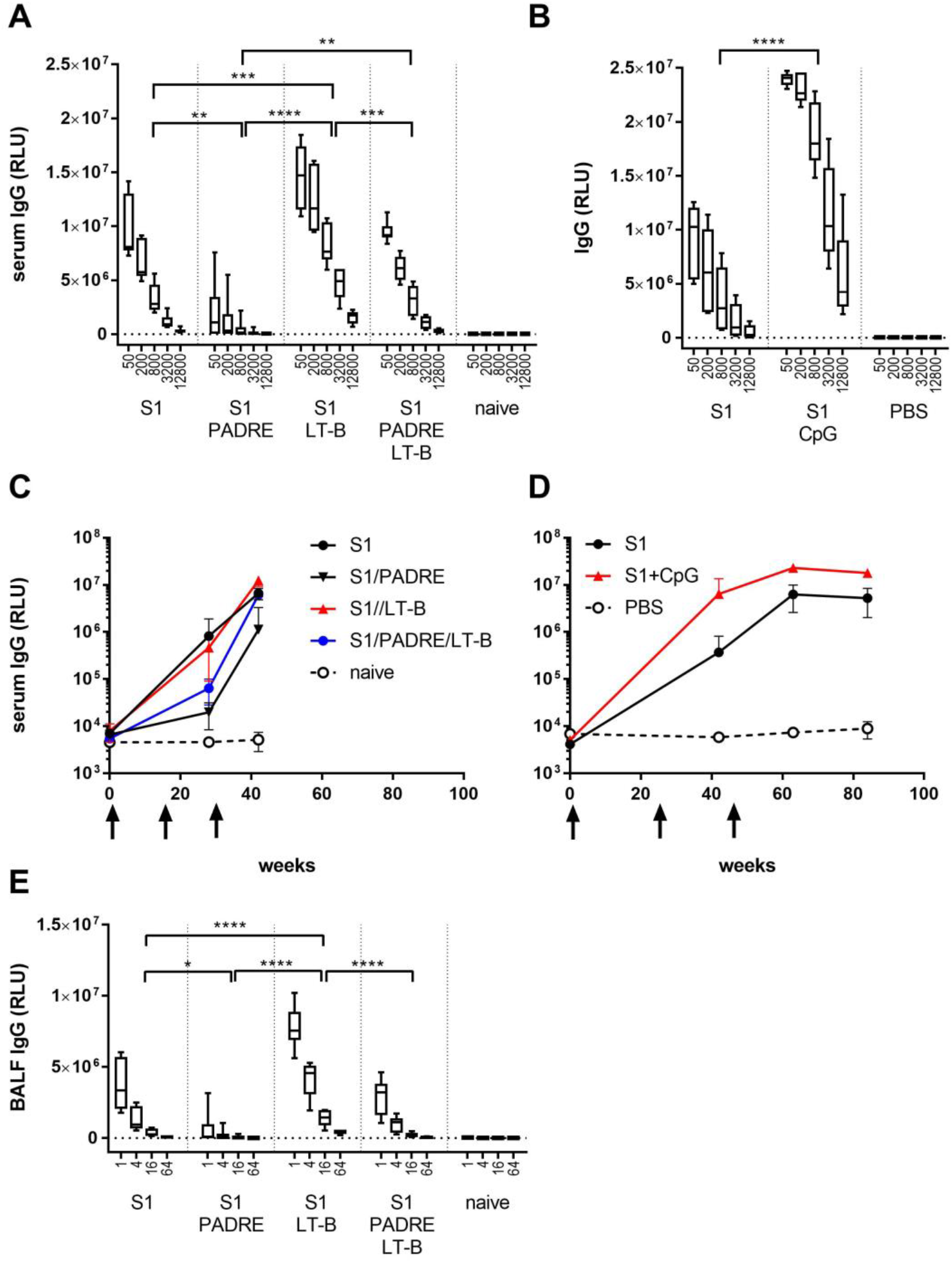
Spike protein-specific IgG in serum and BALF. Mice (n=6) were epicutaneously immunized with recombinant SARS-CoV-2 spike S1 protein with or without different adjuvants (PADRE, LT-B, CpG). Control groups were naïve mice or mice sham immunized with PBS (n=3). Spike protein-specific serum IgG was measured 12 (experiment 1, panel A) or 19 (experiment 2, panel B) days after the third immunization. The time course of the immune response was determined using either 2-week (experiment 1, panel C), or 3-week (experiment 2, panel D) intervals between immunizations. In experiment 1, spike protein-specific IgG was measured in BALF at the end of the experiment (panel E). Data are shown as relative light units (RLU) of a luminometric ELISA (mean±SD). Numbers on the x-axis (panels A, B, E) indicate serum dilutions (1/N). Statistical differences between groups were analyzed by two-way RM ANOVA followed by Tukey’s post hoc test. **** P<0.0001, *** P<0.001, ** P<0.01

Skin immunization with certain adjuvants, such as LT-B, a bacterial heat-labile enterotoxin, has been suggested to promote the induction of mucosal immune responses [19]. Thus, in experiment 1 we investigated, whether epicutaneous immunization with SARS-CoV-2 spike S1 protein (with or without LT-B) would also induce antibody responses in the respiratory tract, the primary affected site in COVID-19 infections. As shown in Fig. 1E, we found elevated levels of spike-specific IgG in the BALF of immunized mice that closely matched the patterns found in serum. However, we could not detect any antigen-specific IgA (data not shown). This might be due to the fact that LT-B is a much less potent mucosal adjuvant than the corresponding (detoxified) holotoxin [20], which was not available for this study.

To investigate whether the antibodies detected by ELISA would also inhibit the binding of ACE2 to spike S1 protein, a SARS-CoV-2 Spike:ACE2 inhibitor assay was performed. This assay was previously established as a reliable surrogate for live virus neutralization [21]. Using serially diluted sera from immunized mice, inhibition curves were fitted using the Sigmoidal 4PL algorithm (GraphPad Prism 7.04), and the EC50 concentration (the concentration required to reduce ACE2 binding to 50%) was calculated. As shown in Fig. 2, both LT-B and CpG significantly boosted the titers of ACE2 binding inhibitory antibodies with average EC50 titers of 1:207 and 1:357, respectively. By contrast, mean EC50 titers in the non-adjuvanted group were 1:21 in experiment 1 and 1:82 in experiment 2 (increased due to a high titer of 1:334 in one animal). For the PADRE group, no titers could be reasonably calculated, as no inhibitory capacity was detected in 4 out of 6 animals.

**Fig. 2.**
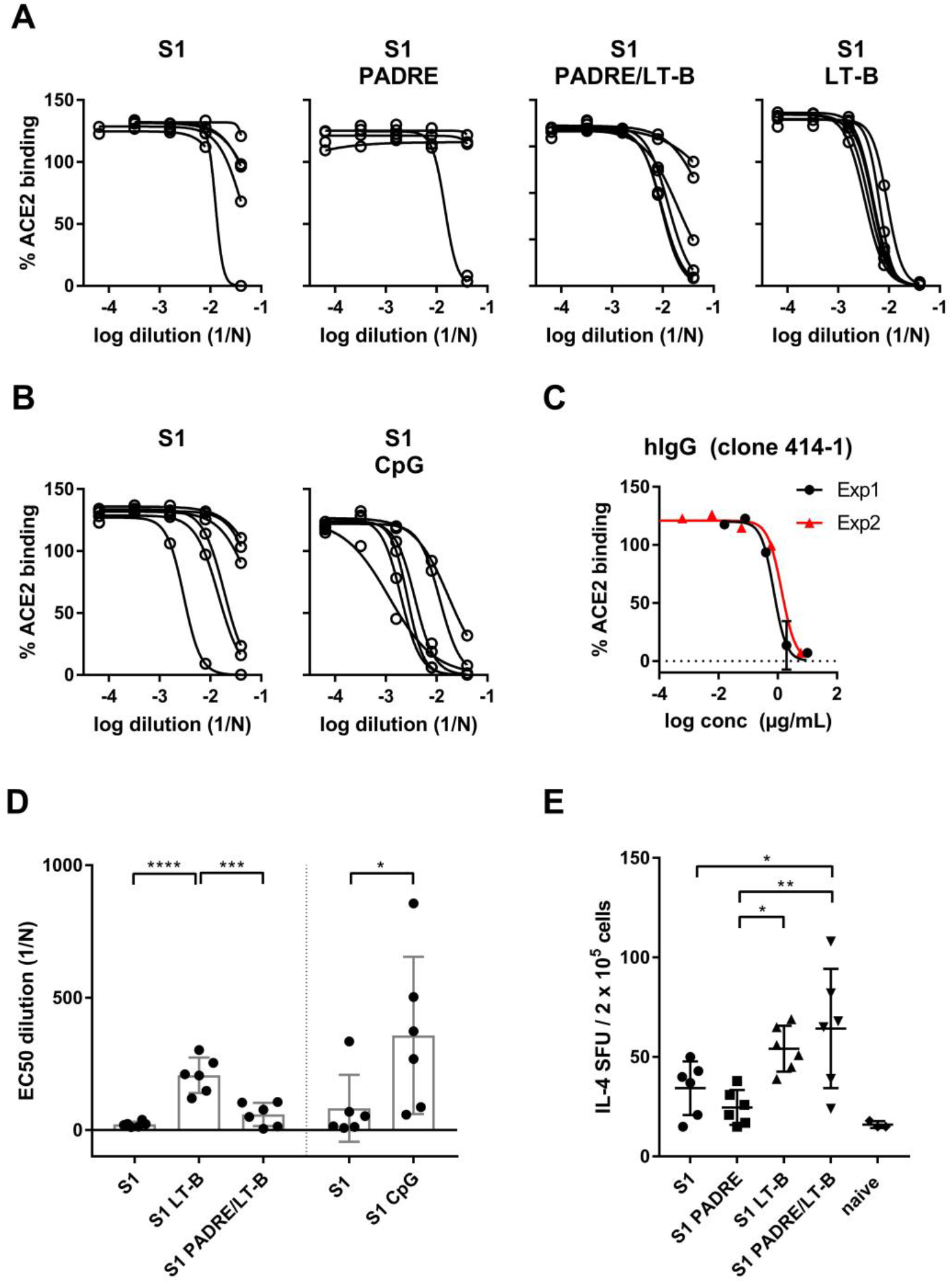
SARS-CoV-2 Spike:ACE2 inhibitor assay and T cell responses. Sera from mice taken after the third immunization were tested for their capacity to inhibit the Spike:ACE2 binding interaction. Inhibition of Spike:ACE2 binding was measured by a luminometric assay and the antibody concentration required to inhibit 50% of Spike:ACE2 binding (EC50) was assessed using fitted inhibition curves. Inhibition curves for individual sera are shown for the groups from experiment 1 (A) and 2 (B). A neutralizing, commercial human IgG antibody (clone 414-1) was used as a reference (C). EC50 values (mean±SD) are shown (D). Splenocytes from immunized mice from experiment 1 were restimulated with spike S1 protein and the number of IL-4 secreting cells (spot forming units – SFU) was assessed by ELISPOT (E). Statistical differences between groups within the individual experiments were calculated by one-way ANOVA and Tukey’s post hoc test (experiment 1), or unpaired T-test (experiment 2). **** P<0.0001, *** P<0.001, * P<0.05.

Although neutralizing antibodies are the main mechanism of preventive antiviral vaccines, the relevance of SARS-CoV-2-specific T cells for protective immunity has been intensely discussed [22, 23]. Thus, in experiment 1, we investigated whether we could detect spike protein-specific TH1/TH2 cells in the spleens of immunized mice. After *in vitro* restimulation with spike S1 protein, we mainly detected IL-4 producing (TH2) T cells. LT-B significantly boosted this T cell response (Fig. 2E). The number of IFN-γ producing TH1 cells was very low, but again, the response could be boosted by LT-B (data not shown).

## Discussion and Conclusions

Laser-assisted epicutaneous immunization with SARS-CoV-2 spike S1 protein induces significant levels of specific serum IgG with ELISA titers of ~1:3200 in mice. Titers can be boosted roughly 5-fold by the use of LT-B and more than 10-fold by CpG-ODN 1826. Elicited serum antibodies are capable of inhibiting the Spike:ACE2 receptor interaction and can thus be expected to be virus-neutralizing [21]. Following immunization with different adjuvants, sera at dilutions of ~1:200 (LT-B) and 1:300 (CpG) showed the same inhibitory capacity as a potent neutralizing human IgG monoclonal antibody (414-1) [18] at a concentration of ~1μg/mL. At this dilution, sera contain roughly 20-40μg of total IgG. Considering that (i) only a minor part of total serum IgG will be antigen-specific and that (ii) hIgG 414-1 was identified as the most potent neutralizing antibody from a convalescent patient [18], we cautiously speculate that these inhibitory titers would probably be protective against infection with SARS-CoV-2. We detected significant IgG, but no IgA antibody responses in BALF. Notably, following natural infection with respiratory viruses, in contrast to the upper respiratory tract, the lung is considered to be mainly protected by IgG transported across the mucosa, and not by secretory IgA [1]. The immunization intervals tested (2 and 3 weeks) had no influence on the magnitude of the antibody response against spike S1 protein; therefore, a rapid immunization protocol might be warranted. Besides the induction of ACE2 inhibitory antibodies, laser-assisted epicutaneous immunization elicited spike-specific T-helper cells possibly contributing to long-term memory responses [22, 23]. Epicutaneous immunization with recombinant spike S1 protein may therefore represent a promising vaccination approach against SARS-CoV-2.

